# Independent evolution of transcript abundance and gene regulatory dynamics

**DOI:** 10.1101/2020.01.22.915033

**Authors:** Gat Krieger, Offir Lupo, Avraham A. Levy, Naama Barkai

**Affiliations:** Department of Molecular Genetics, Weizmann Institute of Science, Rehovot 76100, Israel; Department of Plant and Environmental Sciences, Weizmann Institute of Science, Rehovot 76100, Israel

## Abstract

Changes in gene expression drive novel phenotypes, raising interest in how gene expression evolves. In contrast to the static genome, cells regulate gene expression to accommodate changing conditions. Previous comparative studies focused on specific conditions, describing inter-species variation in expression levels, but providing limited information about variations in gene regulation. To close this gap, we profiled gene expression of related yeast species in hundreds of conditions, and used co-expression analysis to distinguish variations in transcription regulation from variations in expression levels or environmental perception. The majority of genes whose expression varied between the species maintained a conserved transcriptional regulation. Profiling the interspecific hybrid provided insights into the basis of variations, showed that *trans*-varying alleles interact dominantly, and revealed complementation of *cis*-variations by variations in *trans*. Our data suggests that gene expression diverges primarily through changes in promoter strength that do not alter gene positioning within the transcription network.

## Introduction

New phenotypes emerge from mutations that change protein function or protein expression. Between closely related species, protein sequences remain highly conserved while gene regulatory regions vary, suggesting that changes in gene expression play key roles in early evolutionary divergence (King and Wilson, 1975). In support of that, modifying the expression of a single, differentiation-promoting gene during organism development can give rise to novel body morphologies (Prud’homme et al., 2006; Warren et al., 1994). Genomic studies revealed that closely related species differ massively in orthologous gene expression, emphasizing the susceptibility of gene expression for rapid divergence (Combs et al., 2018; Gilad et al., 2006; Goncalves et al., 2012; Landry et al., 2005; McManus et al., 2010; Shi et al., 2012; Tirosh et al., 2006, 2009; Wittkopp et al., 2004, 2008; Yanai and Hunter, 2009; Yvert et al., 2003).

A hallmark of gene expression is its ability to adapt to changing demands. Widespread changes in gene expression that occur during cell differentiation and adaptation to changing conditions are fundamental for these processes. Accordingly, also in the context of the single genome, expression levels measured at one condition provide limited information about expression levels at other conditions. Further, determinants of absolute expression are often different from determinants of gene regulation: the former depends on the efficiency of the general machinery, which is influenced, for example by the presence of a TATA box or the organization of nucleosome along the promoter (Basehoar et al., 2004; Tirosh and Barkai, 2008), while the latter depends on the binding of specific transcription factors (TFs) to gene promoter. Thus, genes may be expressed at similar levels under most conditions but still be subject to different regulation, or, conversely, be subject to identical regulation yet expressed at very different levels.

The ability to control independently expression level and expression dynamics implies that evolution can work on these two properties independently. Distinguishing these two axes of divergence is fundamental for understanding mechanisms of evolution. We reasoned that analysis of regulatory evolution, on a genomic scale, requires a new type of comparative dataset, which goes beyond available comparisons of individual conditions, but survey a wide range of expression profiles obtained under a variety of conditions. Indeed, co-analyzing the expression of different genes across a large number of conditions could distinguish variations in absolute expression from variations in transcription regulation. In addition, such analysis also controls for inter-species differences in environmental sensing or signaling mechanisms, which act upstream of the transcription network. Following this reasoning, we have profiled gene expression of related budding yeast species and their interspecific hybrid under hundreds of conditions, and devised computational approaches for distinguishing variations in expression levels from variations in expression dynamics. We find that massive variations in gene expression levels, coupled with differential environmental perception, mask a largely conserved transcription regulatory network.

## Results

### A compendium of comparative transcription profiles

The budding yeast *S. cerevisiae* and *S. paradoxus* emerged as a model for studying inter-species variations in gene expression (Emerson et al., 2010; McManus et al., 2014; Metzger et al., 2016; Tirosh et al., 2009; Weiss et al., 2018; Yue et al., 2017). The two species diverged ≈5 million years ago, express the same set of genes at a conserved synteny and show 90% and 80% sequence identities at coding and non-coding regions, respectively (Scannell et al., 2011). Further, the two species can readily mate to form viable hybrids. In the context of comparative analysis, hybrids provide a key tool for classifying inter-species variations: the two hybrid alleles are subject to the same *trans* environment so that differences in their expression must result from gene-linked differences (*cis*-effects). The differences between the parents that are lost in the hybrid, are attributed to variations in upstream, diffusible factors (*trans*-effects) (Combs et al., 2018; Dori-Bachash et al., 2011; Emerson et al., 2010; Goncalves et al., 2012; McManus et al., 2010, 2014; Metzger et al., 2016; Shi et al., 2012; Tirosh et al., 2009, 2010; Wittkopp et al., 2004, 2008).

Previous studies described inter-species variations in gene expression when compared at individual conditions (Combs et al., 2018; Gilad et al., 2006; Goncalves et al., 2012; Landry et al., 2005; McManus et al., 2010; Shi et al., 2012; Tirosh et al., 2006, 2009; Wittkopp et al., 2004, 2008; Yanai and Hunter, 2009). We extended the analysis by generating a large collection of comparative transcription profiles surveying a wide array of conditions. Assayed conditions include response to different starvation media, time-resolved progression through the cell cycle, and deletion of several dozens of transcription factors profiled in rich or starvation media (Figure 1A, Table S1). Inter-specific hybrid was generated and profiled under the same conditions. In total, we generated ≈500 transcription profiles for each of the two species and for the hybrid.

Consistent with previous reports, a considerable number of genes differed in abundance between the two species at each individual condition (Figure 1B). A large fraction of these variations remained in the hybrid, indicating their origin in *cis*-acting mutations. On average, in each condition, 1,200 genes showed 2-folds variations between the hybrid alleles (*cis*-effect) and an equivalent number showed same-magnitude *trans*-effects. Compensating and reinforcing *cis*-trans interactions were equally likely and relatively infrequent (Figure S 1A).

Some variations were consistent across conditions, while other were specific to a subset of conditions only (Figure 1C). Consistent with previous reports (Aguet et al., 2017; McManus et al., 2010; Smith and Kruglyak, 2008), *cis* effects were more reproducible across conditions (Figure 1D, Figure S 1C). Both cis and trans effects were enriched with TATA-containing genes (Figure S 1D).

**Figure 1:**
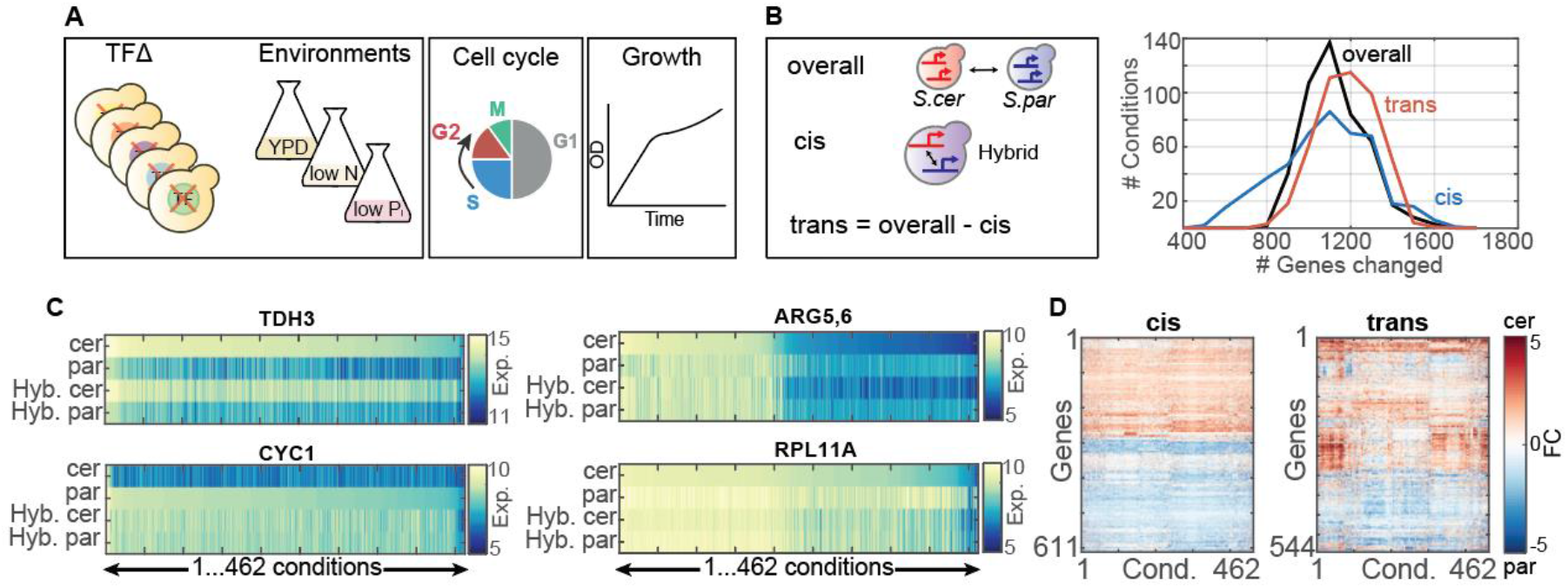
Generating a compendium of comparative transcription profiles. *A. Experimental scheme:* the transcription profiles of S. cerevisiae, S. paradoxus and the interspecific hybrid were measured under the same diverse sets of conditions. *B. Hundreds of genes vary in expression between species in each individual condition:* For each gene, *cis*-effect was defined as the expression difference between the two hybrid alleles, while trans-effect was defined by subtracting the *cis*-effect from the overall between-species variation. For each condition, we counted the number of genes whose expression vary >2 folds between the species (overall), and the number of genes showing >2 folds *cis*- or *trans*-effects. Fold changes were standardize to absolute expression level, shown are Z-score > 1. Shown are the respective distributions, as indicated. *C. Examples of cis and trans varying genes:* Shown are the (log_2_) expression levels of the indicated genes within the indicated species or hybrid alleles, as measured in all conditions in our dataset. Conditions were sorted according to expression levels in S. cerevisiae in *TDH3, ARG5,6* and *RPL11A,* and by S. paradoxus in *CYC1.* *D. Consistency of cis and trans effects across conditions:* Shown are (log_2_) fold changes in cis (left panel) and trans (right panel) of genes with the highest *cis* or *trans* effects in individual conditions (Z-score > 2.5 in at least 10 conditions).

### Quantifying species-similarity in gene regulatory dynamics by comparing co-expression patterns

To compare gene regulation between the two species, we considered two measures of regulatory similarity (Figure 2A). First, we directly compared expression of orthologues across all conditions. This was enabled by our dataset which profiled the two species under the same set of conditions (R^E^, Figure 2A, left). This measure of similarity is sensitive not only to differences in transcriptional regulation, but also to differences in environmental perception or signaling processes acting upstream of the transcriptional network. Second, we compared the pattern at which each orthologue is co-expressed with all other genes in the genome (R^C^, Figure 2A, right). This second measure reports on the relative positioning of a gene within the transcriptional network and is often preferred when analyzing co-regulation within a given genome (Hughes et al., 2000; Ihmels et al., 2005a; Li et al., 2017; Segal et al., 2003), since it is less sensitive to random noise affecting the two genes being compared. Of note, this correlation-based method is less sensitive to variations in environmental perception, but might be biased by large modules of co-expressed genes.

To compare these two measures of similarity, we first considered the ribosomal protein-coding genes, which both species co-induce during rapid growth. The median expression of this group received high similarity score by both measures (Figure 2B), although co-expression similarity was somewhat higher (0.86 vs. 0.76). A clear difference between the two measures, however, appeared when comparing individual genes: expression-based similarities (R^E^) were widespread and often low, while correlation-based ones (R^C^) were consistently high (Figure 2C).

Extending the analysis to all orthologues, we find expression-based similarities (R^E^) are relatively low and widespread while correlation-based ones (R^C^) are significantly higher and tightly distributed (Figure 2D). As a control, we applied these same measures to define similarities within each individual genome. First, we considered a nearest-neighbor (NN) control, whereby each gene is assigned a NN, namely the gene to which it is most similar (Figure 2E). Also here, correlation-based NN similarities (R^C^) were higher and more tightly distributed compared to expression-based ones (R^E^) (Figure S 2E). Next, we considered a dataset-control, whereby we split the datasets into two random parts and examine whether co-expression values assigned to each gene remain invariant. The majority of genes obtained high R^C^ similarity score in this dataset control (Figure 2E), and in analogous comparison with external datasets (Figure S 2A-D), confirming relative invariance of this measure to the precise dataset used. Together, we conclude that the correlation-based score, R^C^, is suitable for capturing regulatory similarities and we therefore proceeded with this measure.

Using co-expression (R^C^) as our measure of regulatory similarity, about half of orthologues were assigned high similarity scores in the range 0.8-0.9 (Figure 2D, E). The absence of values higher than 0.9 suggests a systematic bias caused by diverging genes. Indeed, for 552 orthologues, similarity scores were lower than 0.5. By comparison, NNs or dataset-control similarities were mostly at the range of 0.95-1, with only a negligible fraction being < 0.5 (Figure 2E). Together, this suggest that while the majority of orthologues maintained a conserved regulation, regulatory variations can still be detected in a significant number genes.

**Figure 2:**
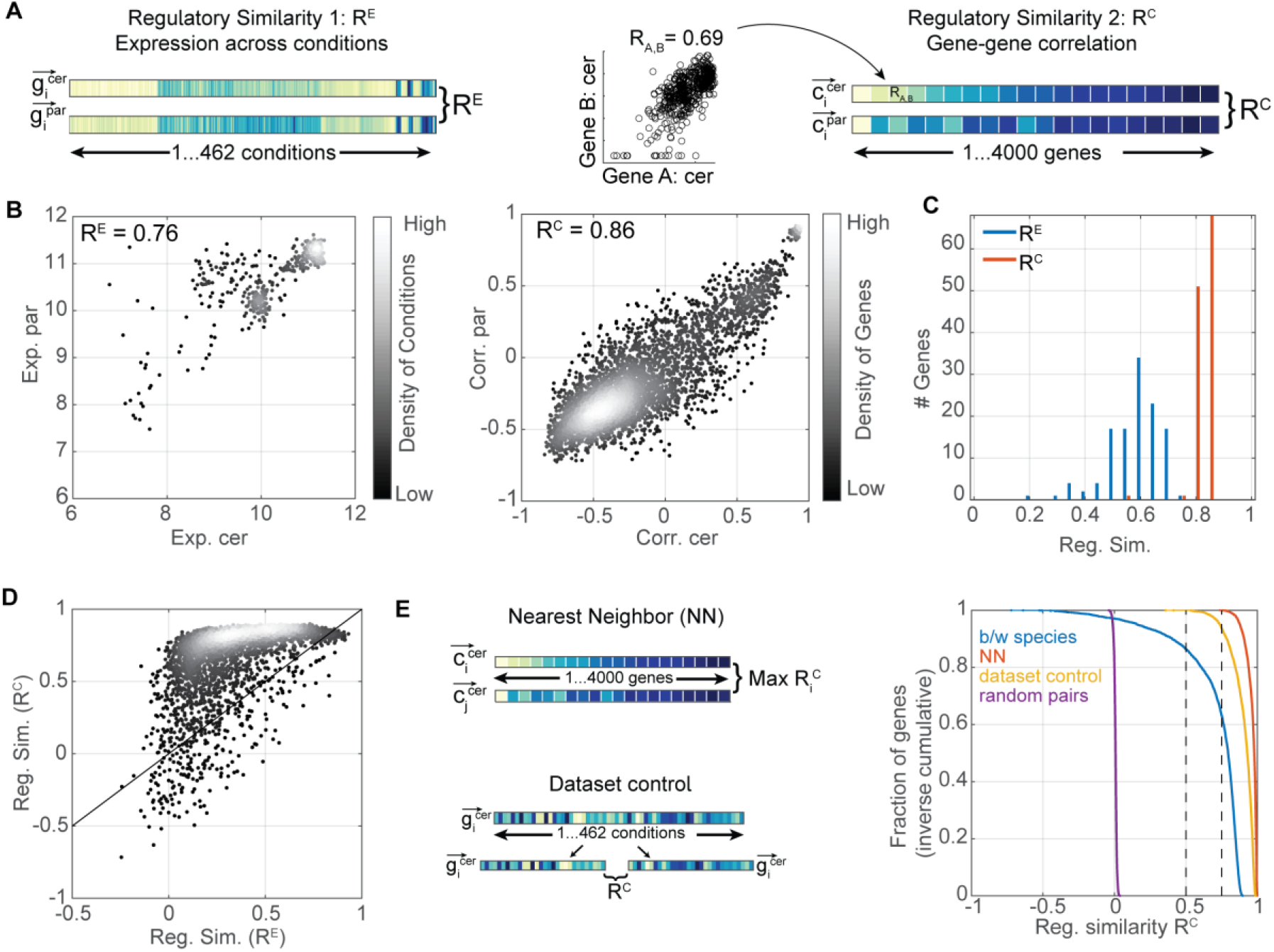
Gene co-expression as a measure of regulatory similarity. *A. Measures for regulatory similarity, a scheme:* Regulatory similarity can be measured by directly correlating expression of orthologous genes over the set of measured conditions (R^E^, left) or by correlating the pattern of co-expression with all other genes in the genome (R^C^, right). *B. -C. Regulatory similarity of ribosome-coding genes:* Shown is the median (log2) expression of 152 ribosome-coding genes in S. cerevisiae and S. paradoxus at each tested condition (left), and the co-expression of this median with every gene in the genome (right). Note that both measure suggest high regulatory similarity (R^E^ = 0.76, R^C^ = 0.81). The distribution of regulatory similarities assigned to individual orthologues is shown in C. *D. Correlation-based regulatory similarities (R^C^) are higher and more tightly distributed compared to expression-based similarities (R^E^*): shown are the respective similarity scores for all orthologous. *E. Regulatory similarity between orthologues are comparable to these found within individual genomes:* Two within-genome controls were defined, as shown. First, Nearest neighbors (NN) are defined by pairing each gene with its most similar partner. Second, dataset control is obtained by splitting the set of conditions into two random subsets and measuring the gene regulatory similarity R^C^ between the two datasets. Shown are the distributions of Regulatory similarity scores.

### Regulatory variations are distinct from variations in expression levels

We asked whether genes that vary in expression levels also show low levels of regulatory similarity. Examining individual genes suggested to us that these two measures are distinct; Stress-induced genes, for example, varied in absolute expression across many conditions, yet their regulatory similarity was high, whereas genes coding for the mitochondrial ribosomal proteins were expressed at similar levels by the two species, yet their regulatory similarity was low (Figure S 3A). Comparing the regulatory similarity of each gene with the extent to which its expression levels varied across all conditions, we saw no consistent correlation between these two measures (Figure 3A). Further, also when testing individual conditions, we find no enrichment of regulatory variations within the top 100 *cis* effects and only a moderate enrichment within the top 100 *trans* effects (Figure 3B). Together, we conclude that regulatory variations, as captured by co-expression analysis, are distinct from variations in absolute expression.

To begin classifying genes of low regulatory similarity, we considered the allele-specific expression within the hybrid. These two alleles are subject to the same *trans* environment and, accordingly, are typically more similar than the respective orthologues (Figure 3C). We also noted that of the 552 orthologues whose similarity score was <0.5, 259 maintained a high similarity between the hybrid alleles (>0.75), and we classified these as pure regulatory *trans* effects (Figure 3C). Still, in 31 cases, similarity between the hybrid alleles was lower than 0.5 and these were classified as regulatory *cis* effects (Figure 3E).

Having this classification of regulatory *cis* and *trans* effects, we asked whether these effects show some tendency for increased variation in absolute expression. To this end, we first considered the median *cis* or *trans* variations across all conditions (Figure 3, D,F, top panels), and examined variations observed in individual conditions (Figure 3, D,F, bottom panels). Overall, regulatory *trans* effects were biased towards a lower, rather than higher variation in absolute expression (Figure 3D). Regulatory *cis* effects did show some tendency for increased *cis*-(but not *trans*-) variations in absolute expression, but this tendency was relatively minor. We conclude that, in large, regulatory variations captured by our correlation-based measure of similarity are distinct from variations in absolute expression.

**Figure 3:**
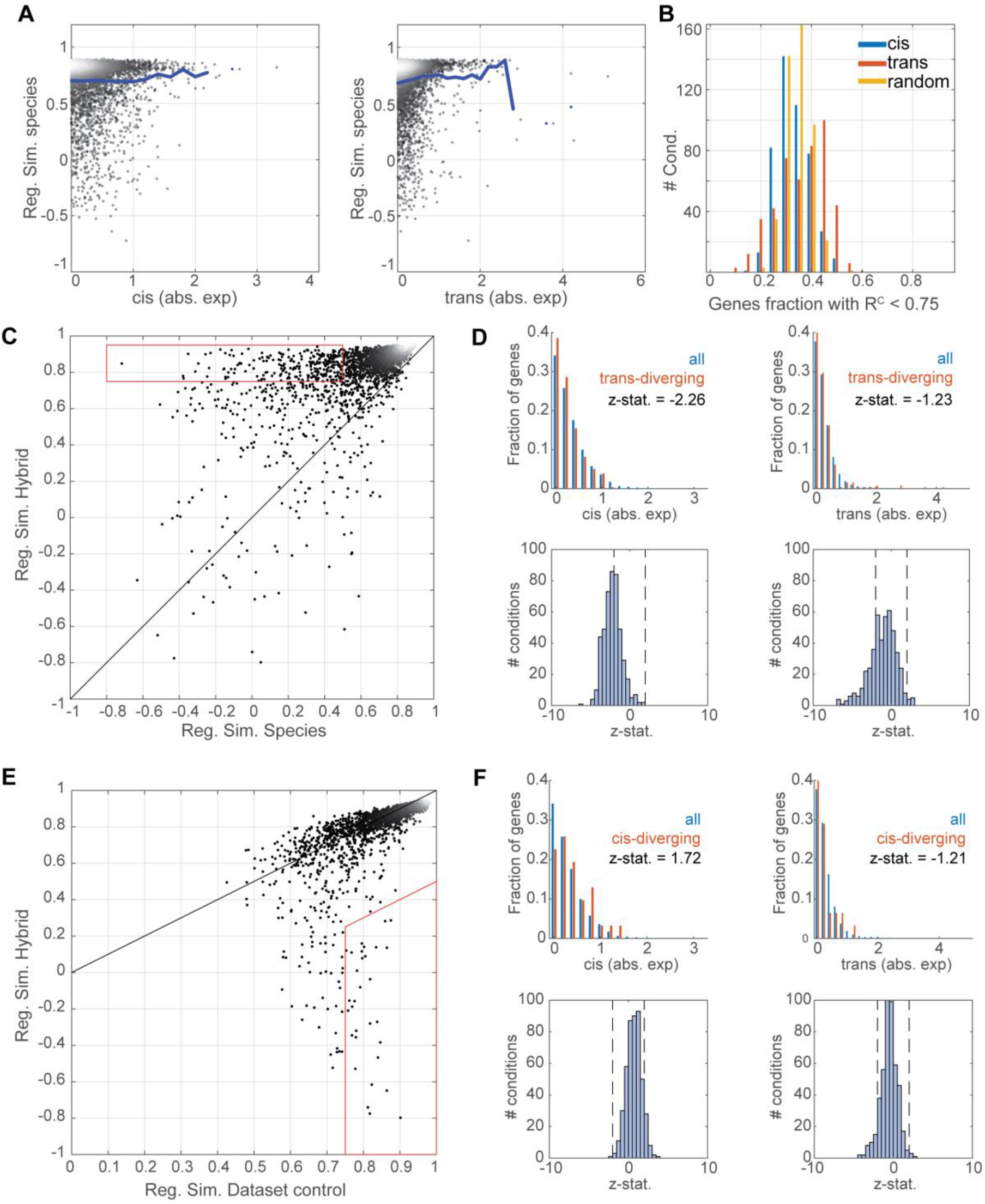
Regulatory variations are independent from variations in expression levels. *A-B: Regulatory similarity correlates poorly with variations in expression levels:* shown in (A) is the regulatory similarity (R^C^) between orthologues as a function of the *cis* or *trans* variations in absolute expression. Here, we considered the median variation in absolute expression, as measured across all conditions. Line represent the mean regulatory similarity at the indicated *cis-* or *trans-* effect (left and right panels, respectively). In (B) we considered the top 100 *cis* and *trans* variations in absolute expression at each individual condition, and measured the number of genes showing low regulatory similarity (<0.75). Shown are the respective distribution, compared to random control. Note that *cis* effects are depleted of low-regulatory similarities, while *trans* effects show higher fraction of low-regulatory similarity (p-value = 10^-11^, Kruskal-Wallis test). This analysis was repeated for different regulatory similarity thresholds as shown in Figure S 3B. *C-D: Regulatory trans-variations are depleted of absolute expression-variations:* regulatory trans-variations were defined as these showing low (R^C^<0.5) similarity between species, while maintaining high (R^C^>0.75) similarity between hybrid alleles, as shown in (C). The distribution of expression-variations within this group of 259 genes was than examined in (D). This was done first by considering the condition-median *cis* or *trans* effects (top panels). Next, the respective distributions were analyzed for each condition separately, and are summarized by the significance of their deviation from the control all-genes distribution (z-statistics). Note that a negative z-statistics indicate that the set of *trans* varying genes is depleted of the respective *cis* or *trans* variations in absolute expression. E-F. *Regulatory *cis*-variations show a limited tendency for absolute expression-variations:* same as C-D above, but for the group of regulatory *cis*-variations. Genes in this group show low regulatory similarity between hybrid alleles, while maintaining high (>0.75) regulatory similarity in the mean dataset control of the two alleles.

### Patterns of trans-dependent regulatory variations

Considering the prominence of *trans* effects among genes of low regulatory similarity, we decided to first focus on *trans* varying genes. Illustrative examples include VTC3, a vacuolar transporter chaperone, and PMT2, a protein involved in ER membrane control. In both cases, orthologues show low regulatory similarity while the hybrid alleles remain highly similar (Figure 4A, B). Further, in both cases, the hybrid alleles were highly similar to one of the parents. Other properties, however, differed between these two cases. VTC3 maintained its high correlation with phosphate-responding genes in both species (Figure 4A, Figure S 4A), suggesting that its direct regulator, Pho4, is still active, but differentially regulated in the two species. Indeed, Pho4 deletion affected VTC3 expression in a condition- and species-specific manner. PMT2, on the other hand, have lost signs of co-regulation in S. paradoxus: its correlation with all other genes is low, and is only moderately reproducible in our dataset control (R^C^= 0.77, 17^th^ percentile). Therefore, it is likely that the TF regulating PMT2 in S. cerevisiae and the hybrid shows a limited, or lack of activity in S. paradoxus.

The high similarity of the hybrid alleles with one of the parents, observed in both examples, was surprising to us, since we expected that combining differentially regulated *trans*-acting factors within the same genome would result in additive or synergistic effects. Examining the full set of *trans* diverging genes we found dominance to be general: in the majority of cases (210/259), the hybrid alleles were clearly paradoxus-like or cerevisiae-like (Figure 4C). This widespread dominance suggests that allelic-variants are compensated when combined in the hybrid.

A case where dominance is expected is of a TF that becomes inactive in one species. This may result from an inactivating mutation, or from a regulatory change that affects its activity in the conditions we surveyed. In this case, the hybrid alleles will be similar to the active species. Nearest-neighbor (NN) similarities may be indicative of such cases, as these commonly arise from regulation by the direct TF and will be lost upon its inactivation. Indeed, NN similarities <0.85 were found for 40% of *trans* diverging genes, compared to 8% of all genes (Figure 4D, Figure S 5B). Further, in the vast majority (94-98%) of cases, the dominant species maintained high NN correlation. Therefore, these cases of low NN similarity in the recessive species likely result from loss of activity of the direct regulator.

We next considered cases of high NN similarity. Here, variation could result from differential regulation of the same direct TF, as exemplified by VTC3. Alternatively, the dominant regulatory TF may differ between the species. To distinguish between these possibilities we asked whether NN similarities are maintained across species (Figure 4E). As expected, NN relations were conserved in the case VTC3, as well as in other Pho4-regulated genes. Other groups showing this behavior were associated with functions such as iron transport and fatty-acid synthesis in the S. paradoxus dominance class, or mitochondrial translation and cell wall maintenance in the S. cerevisiae dominance class (Figure 4F). By contrast, in other genes, NN relations were maintained in only the dominant, but not in the recessive species, or were lost altogether. In these cases, the recessive species may have lost the primary regulation acting in the dominant species and in the hybrid, as in PMT2. We conclude that *trans*-dependent regulatory divergence result from differential regulation of specific TFs in the two species, with a considerable number of cases corresponding to the lack, or limited TF activity in one of the species.

**Figure 4:**
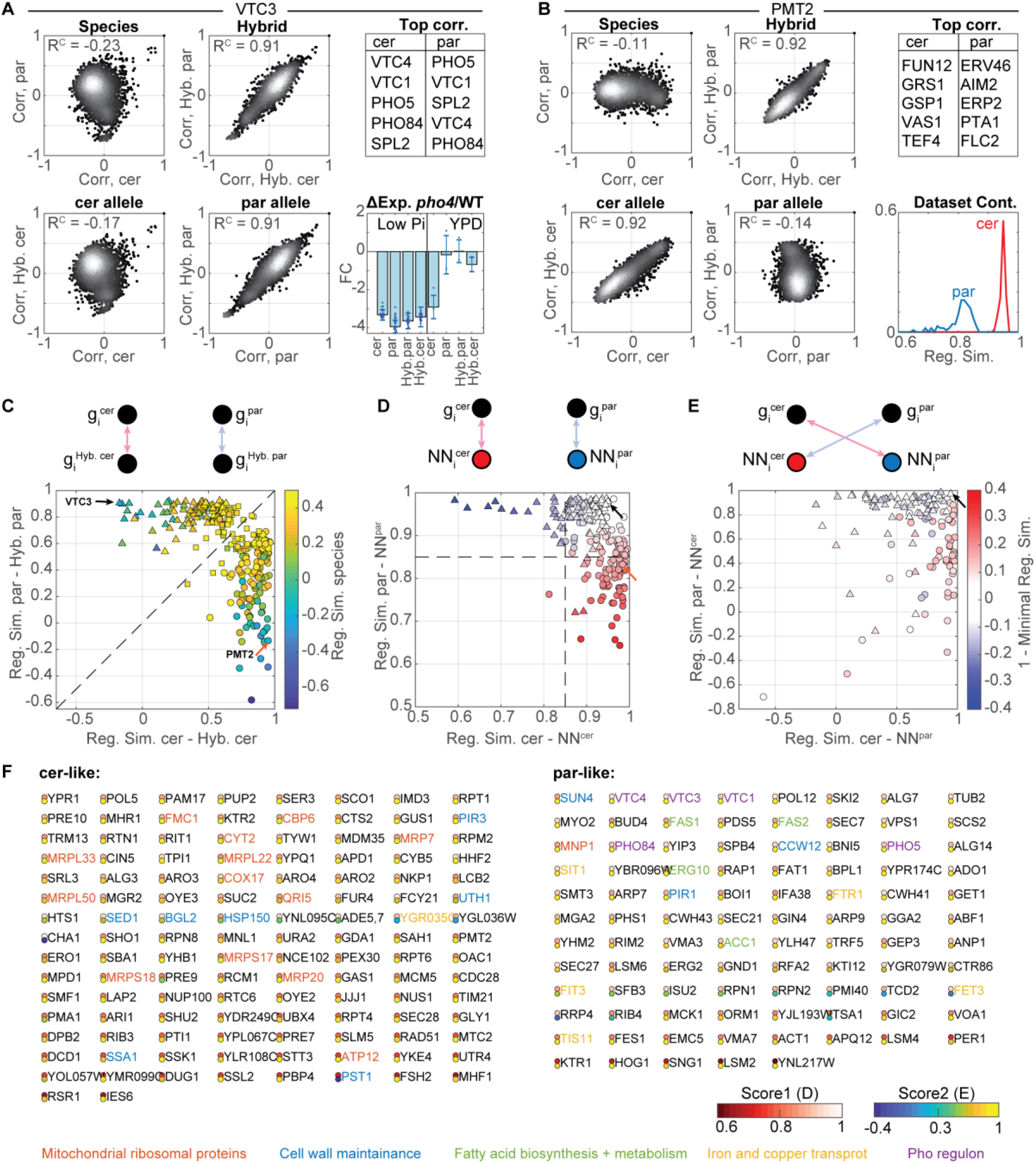
trans-dependent regulatory variations show dominant effects. *A-B: Examples of regulatory trans-variations:* The four panels on the left compared the co-expression of VTC3 (A) or PMT2 (B) with each gene in the indicated genomes. Each dot is a gene and color-code indicate gene density. Also shown are the highest-correlated genes in each species (right, top), the change in VTC3 expression upon PHO4 deletion (A, bottom right) and the distributions of PMT2 regulatory similarity values in obtained in 100 repeats of dataset controls. *C. Hybrid alleles show high similarity to one of the parents:* Shown is the regulatory similarity between the hybrid allele and the corresponding parent. Color-code indicates similarity between the parents. Triangle and circles indicate paradoxus-like or cerevisiae-like dominance, respectively; Squares indicate genes equally similar to both species. *D. Nearest-neighbor (NN) similarities are higher in the dominant species:* Shown is the regulatory similarity of each trans-diverging gene to its NN in the two species, as indicated in the scheme. Color-code is based on the lower NN similarity, taken as positive when S. cerevisiae is lower and negative when it comes from paradoxus. Dot shape as in C. *E. Cross-species NN similarities vary between genes:* Shown are cross-species NN similarities, as indicated by the scheme at the top. Dot shape as in C, color-code as in D. *F. Regulatory trans-variations:* Shown are the genes in the regulatory *trans*-variations, classified by the dominant species and ordered by the two regulatory scores corresponding to their positions in Fig. D-E, see methods for details.

### Patterns of *cis*-dependent regulatory divergence

Next, we examined the class of regulatory *cis*-variations. Illustrative examples are LSO2 and PGM1. In *S. cerevisiae*, LSO2 is annotated as ribosome-associated gene (Figure 5A). Examining the LSO2 promoter, we noted that S. cerevisiae have gained a binding site for SFP1, the principle TF regulating ribosomal protein and ribosomal biogenesis genes. Consistently, LSO2 was tightly co-expressed with genes coding for ribosomal proteins in S. cerevisiae but not in S. paradoxus. Phosphoglucomutase 1 (PGM1), as well as other glycolytic genes, is known to be regulated by the TF GCR1 in S. cerevisiae (Nishi et al., 1995). Its co-expression pattern varied in cis, being coexpressed with other glycolytic genes in S. cerevisiae but not in S. paradoxus. This is consistent with the presence of the GCR1 binding motif in S. cerevisiae but not in S. paradoxus (Figure 5B). These two cases may therefore exemplify regulatory variations resulting from a gain of TF binding.

We noted that in terms of regulatory similarity, the S. cerevisiae orthologue of LSO2 is highly similar to the respective hybrid allele (R^C^=0.87). Examining the full set of regulatory *cis*-variations revealed this to be general (Figure 5C). In fact, for the most pronounced variations, both hybrid alleles were similar to their respective parents, suggesting distinct regulation of the two alleles by different TFs. By contrast, moderate correlations between hybrid alleles were often associated with low similarity with at least one of the parents (Figure 5C) and correspondingly low NN similarities (Figure 5D), suggesting limited regulation of the respective allele. Consistent with this possibility, *cis*-diverging genes more often changed their gene-neighborhood compared to *trans*-diverging genes (Figure 5E). As expected we observe no enrichment for common functions among the *cis* varying genes (Figure 5F). Taken together, our results suggest that the majority of *cis* effects result from differential regulation of the two alleles, either by two distinct TFs, or due to a loss or gain of TF regulation on one of the alleles.

The low regulatory similarity between LSO2 hybrid alleles was somewhat suppressed when comparing the parental species, indicating a compensating effect acting in *trans*. Examining the full set of *cis*-effects, we found *trans*-effects to be prominent among genes showing *cis* variations (Figure 5G). While only ~20% of genes showed species-hybrid difference >0.2, 70% of the *cis*-varying genes showed such an effect. Notably, those *trans* effects were biased towards compensating, that is, decreasing the difference between species relative to the hybrid alleles. We conclude that *cis*-dependent regulatory variations are often accompanied by additional, often compensating, *trans* effects.

**Figure 5:**
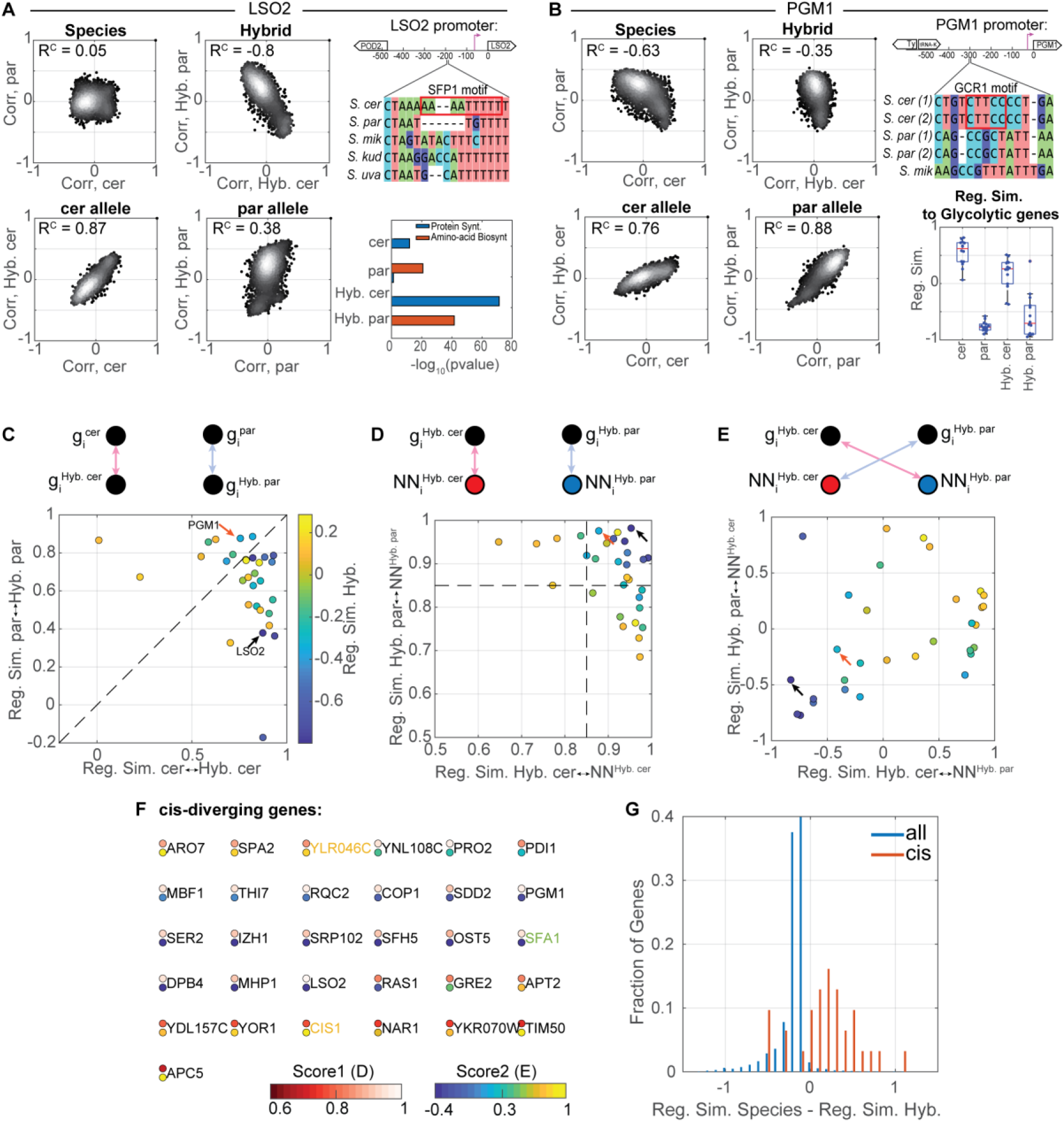
cis-dependent regulatory variations are compensated by trans effects. *A-B: Example of regulatory *cis*-variation:* plots on the left are the same as in Fig. 4A-B. Shown on the right is section of S. cerevisiae LSO2 promoter containing the SFP1 motif, and its comparison with the promoters of the indicated species. Enrichment of ribosome-coding genes within the top-100 LSO2 correlating genes in the different genomes is also shown and is compared with the respective enrichment for genes in the amino-acid biosynthesis group. For PGM1 (B), shown on the right is section of S. cerevisiae PGM1 promoter containing the GCR1 motif, and its comparison with the promoters of the indicated species. As this promoter sequences is highly divergent in distant species, we show alignment of two S. cerevisiae strain (S288c, SK1), two S. paradoxus strains (CBS432, N44) and S. mikatae. Regulatory similarities (R^C^) of PGM1 with genes involved in glycolysis and co-expressed with TDH3 are shown for each genome. *C. High regulatory similarity between hybrid alleles and the respective orthologues:* Shown is the regulatory similarity between the hybrid allele and the corresponding parent (as in Figure 4C). Color-code indicates the similarity between hybrid alleles. *D. Nearest-neighbors similarities are high in both hybrid alleles:* Shown are NN similarities of the regulatory *cis*-varying genes, as in Figure 4D. Color-code as in 5C. *E. Cross-alleles NN similarities vary between genes:* Shown are the cross-alleles NN similarities, as in Figure 4E. Color-code as in 5C. *F. List of *cis*-diverging genes:* regulatory scores and functional gene groups as in Figure 4F. *G. regulatory *cis*-variations are often accompanied with regulatory trans-variations:* For each gene, we calculated the difference in the regulatory similarity between orthologues and the regulatory similarity of the respective hybrid alleles. Shown are the distributions of these scores for the set of *cis*-varying genes, and for the full genome.

## Discussion

Understanding how gene expression evolves is a major challenge. In this work, we distinguished two properties of gene expression: transcript abundance and dynamic regulation. It is a common observation that genes that are subject to a similar regulation can still be expressed at widely different levels. This ability to control independently the level and regulation of gene transcription implies that both can be subject to evolution.

Previous studies that compared gene expression between related species considered a few conditions and therefore could only address variations in absolute expression levels. Whether these variations result from changing regulation or from differences in expression levels remained unresolved. By generating a compendium of comparative data, in which we surveyed a diverse set of conditions, we distinguished variations in expression level from variations in dynamic regulation. Our major finding is that these two variation types are distinct. In particular, the majority of genes whose abundance vary between the species, did maintain a conserved regulatory pattern.

We based our measure of regulatory similarity on the pattern of gene co-expression. This measure is commonly used for defining co-regulation within individual genomes (Hughes et al., 2000; Li et al., 2017; Segal et al., 2003), and was previously used for comparing core-modules in more distant species (Ihmels et al., 2005a, 2005b; Thompson et al., 2013; Tsaparas et al., 2006). It is beneficial in our context as it minimizes differences in environmental perception, which are upstream to the transcription network itself. We note that systematic application of this method for identifying regulatory-varying genes requires close enough species that express largely the same set of genes and where regulatory variations are the exception, rather than the norm.

To gain insights into the mechanistic basis of regulatory variations, we used an interspecific hybrid. A large fraction of regulatory variations observed between the species was lost when comparing the two hybrid alleles. This indicates prominence of *trans* effects, in agreement with other experimental systems (Albert et al., 2018; Liu et al., 2019; McManus et al., 2010; Sanchez et al., 2019; Yvert et al., 2003).

We expected that the *trans* effect will show additive or synergistic interactions within the hybrid. Contrasting this expectation, we found that the vast majority of trans-effects interacted in a dominant way, being practically identical between the hybrid and one of the parent. This may be explained if *trans* divergence results from a limited activity of a TF in one of the species, which indeed appears to explain a considerable fraction of *trans*-effects we observe. This may provide a mechanistic explanation for the phenomenon of genome asymmetry that was reported before in a variety of organisms (Feldman et al., 2012; Lemos et al., 2008; Ren et al., 2019) whereby the phenotypic manifestation of some traits, including gene expression patterns, favors one genome over the other rather than being intermediate.

We measure regulatory similarity using a correlation-based measure, while variations in transcript abundance is measured in units of read counts. It therefore difficult to conclude which of these variations is more prominent. Our analysis of the data, however, suggested to us that cases in which genes have lost or gained regulation by specific TFs are rather rare, and that most variations result from limited activity of a common regulator. Overall, our data suggests that the massive variation in absolute gene abundance masks a remarkably conserved transcription network.

## Materials and methods

### Yeast Strains

Yeast stains in this study were constructed on the background of S. cerevisiae S288c and S. paradoxus CBS432 (OS142) and their hybrid. Strains listed in Table S2. In this study we expression-profiled only diploid yeast cells. transcription factors that were deleted are listed in Table S1. All transformations were done using the standard LiAc/SS-Carrier/PEG transformation method.

### Time course experiment in low nitrogen median for multiple strains

Glycerol stock libraries were plated on YPD-agar plates and re-plated on selective agar plates (YPD+G418+Hyg+Nat) after two days. Colonies were pinned to liquid starters, grown to stationary phase for 24 h, than diluted in 1.5 ml YPD in 2 ml deep-well plates. These plates contained one glass bead per well for proper mixing of the culture. In these experiments, each half plate contained a technical repeat of the same culture; one half was used for optical density (OD_600_) measurements and the other for cell harvesting for RNA purification. The cultures were grown in 30° C shaker for 6-8 h reaching to OD_600_ of 0.1-0.6, than sampled and washed once, resuspended in nitrogen-depleted medium and grown in the same conditions, sampled 1 h and 16 hours post medium shift. Each sample contained 0.5 ml culture; yeast culture was centrifuged at 4000 rpm for 30 seconds, supernatant was removed using multipipette vacuum and pellets were immediately frozen in liquid nitrogen and stored in −80° C.

### Nitrogen-depleted medium

0. 67% Yeast Nitrogen Base without amino acids and ammonium sulfate (Bacto-YNB), 2% D-glucose, 0.05 mM ammonium sulfate, 20 mg/l Urcail, 20 mg/l Histidine, 100 mg/l Leucine.

### Cell cycle time course

Yeast starters were grown in YPD over night at 30°C to stationary phase, and were inoculated to fresh medium to OD_600_ of 0.005 in 100 ml in a 500 ml flask, then grown overnight. When reaching OD_600_ of 0.1-0.2, hydroxyurea (HU) was added to the media to a final concentration of 0.2 M for additional 2 hours. To remove HU from the media, the cells were washed twice by centrifugation (4000 rpm for 1 minute) and re-suspended in fresh, warm, equal-volume YPD. Then, the culture was returned to a bath orbital shaker. Cells were collected at the following time points: before HU, 5’,10’,20’,30’,60’ and 120 minutes in HU, and every 5 minutes after release for 3 hrs. In total 43 time points for each strain. For RNA, samples of 1.5 ml were taken and centrifuged for 10 seconds in 13,000 rpm, sup was removed and the pellets were immediately frozen in liquid nitrogen. For DNA staining (to assess proper synchronization, data not shown), samples of 1.5 ml were taken were taken and centrifuged for 10 seconds in 13,000 rpm and resuspended in cold 70% ethanol. Samples were kept in 4°C. This experiment was carried with two independent biological repeats for each strain.

### Growth curve time course

Overnight starters of yeast were diluted in 50 ml YPD in 250 ml flasks and grown in a bath orbital shaker of 30°C overnight (12 hours). The time course started when the cultures reached OD_600_ of 0.4-0.5 and were sampled for OD_600_ measurements in RNA extraction in the following time points: 80, 125, 170, 215, 260 minutes, 5.1, 6.1, 7.1, 8.1, 9.1, 10.1, 11.1, 23.2, 32.5 and 49.5 hours after start.

### Time course of transition to phosphate starvation

Overnight starters of yeast were diluted in 100 ml SC in 500 ml flasks and grown in a bath orbital shaker of 30°C overnight (12 hours). When the cultures reached OD_600_ of 0.2 they were sampled for RNA extraction, then washed twice in SC medium lacking Pi and inoculated to that media. For the next 6 hours the cultures were sampled every 15 minutes for RNA extraction. At the end of the time course the three cultures (WT strains of S. cerevisiae, S. paradoxus and the hybrid) have completed 1.5 rounds of mitotic divisions. The experiment was repeated in a small scale to include ΔPHO4 strains of the two species and hybrid.

### RNA extraction and sequencing

RNA was extracted using a modified protocol of nucleospin^®^ 96 RNA kit. Specifically, cells lysis was done in a 96 deep-well plate by adding 450 μl of lysis buffer containing 1 M sorbitol (SIGMA-ALDRICH), 100 mM EDTA and 0.45 μl lyticase (10 IU/μl). The plate was incubated in 30°C of 30 minutes in order to break the cell wall, then centrifuged for 10’ at 3000 rpm, and supernatant was removed. From this stage, extraction proceeded as in the protocol of nucleospin^®^ 96 RNA kit, only substituting β-mercaptoethanol with DTT. cDNA was prepared from the RNA extracts, barcoded using Tn5-mediated tagmentation protocol, and sequenced using Illumina NextSeq.

### Library preparation and sequencing

Libraries were prepared as described in Voichek et al., 2018. Poly(A) RNA was selected for by reverse transcription with a barcoded poly(T) primer. The barcoded DNA-RNA hybrids were pooled and fragmented by a hyperactive variant of the Tn5 transposase (courtesy of Ido Amit). Tn5 was stripped off the DNA by treatment with SDS 0.2% followed by SPRI cleanup and the cDNA was amplified and sequenced with Illumina NextSeq 500 with 50 bp reads.

### Dual-genome alignment pipeline

The following pipeline was designed with- and implemented by Gil Hornung (INCPM, WIS). Sequenced reads were mapped against the concatenated genomes of S. cerevisiae and S. paradoxus strain CBS432 (Yue et al., 2017). Mapping was performed with STAR 2.4.2a (Dobin and Gingeras, 2015). The alignments were divided into genomes based on the alignment scores (attribute AS:I in SAM format). Uniquely mapped reads were assigned to the orthologue with the better score. If there is no difference in the scores between the two genomes and the alignment is unique in the cerevisiae genome, then the alignment to the cerevisiae genome is kept and assigned as indistinguishable (same way for the paradoxus genome). Counting was performed on the TES (Transcript End Site) of each gene. The TES was defined between −500 to +200 bases relative to the stop codon. The reads were counted using htseq-count.

### Data filtering and normalization

Each sample was normalized by its sum of mapped reads and multiplied by one million, transformed to log_2_ scale, and values lower than 3.32 were assign to NaN. Samples with less than 150,000 reads were filtered out. Reads of conserved genes that were determined as indistinguishable were assigned to each sample, and were divided equally between the two genomes in the hybrid samples.

### Cis and trans effects on absolute expression

For each gene in each condition we assigned the following measures: interspecies fold change: [log_2_(# reads of cerevisiae) - log_2_(# reads of paradoxus)], *cis:* [log_2_(# reads of cerevisiae in the hybrid) - log_2_(# reads of paradoxus in the hybrid)], *trans:* [interspecies-cis]. For each effect (interspecies, *cis, trans)* we generated and MA plot (fold change vs. product that resembles the gene expression level). The product was divided into 10 bins, for each a z-score of the fold change was computed. Differentially expressed genes depicted in Figure 1 B,D and Figure S 1 are those that exceed fold change of 2 (>=1 in log_2_ scale) and z-score of 1. Figure 1 D presents genes with extreme divergence, hence z-score > 2.5 in at least 10 experimental conditions, for *cis*-effects and for trans-effects separately.

### Enrichment for functional groups

Enrichment analysis was done through applying a hypergeometric test to a list of genes against functional genes groups that include: expression modules (Ihmels et al., 2002), Environmental Stress response (Gasch et al., 2000), GO slim, Transcription factors targets (MacIsaac et al., 2006), KEGG pathways, expression levels, burst size (Newman et al., 2006), TATA-containing promoters (Basehoar et al., 2004), OPN and DPN (Tirosh and Barkai, 2008).

### Correlation across conditions (R^E^)

For each dataset (cerevisiae, paradoxus, hybrid-cerevisiae and hybrid-paradoxus) each gene was assigned with an expression vector spanning all common conditions (462). Expression vector of the i^th^ gene in cerevisiae is therefore: 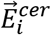, and correlation between species across conditions is: 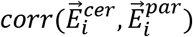. We applied Pearson’s correlation for all correlation computations in this study.

### Regulatory similarity by co-expression (R^C^)

Here, pairwise correlation matrices for all genes in the genome were computed per dataset. To filter out lowly expressed and non-regulated genes, we considered correlation coefficient >|0.21 as significant. Gene with correlation vectors that comprise of more than 90% nonsignificant values, or less than 100 significant values were filtered out, resulting in 4059 genes in the analysis. The inter-species correlation is defined per gene i as: 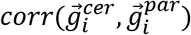. To control for repeatability of the signal, we computed the correlation with two random halves of conditions in each dataset. The median value of 10 permutations of this computation represents repeatability (Figure 2 D). As a second control measure, we computed the correlation of a gene to its nearest neighbor in the correlation-vector space. That by computing the correlation of a pairwise-correlation matrix with itself, and present the maximum per gene.

### Regulation scores

To define whether a trans-diverging gene lost or maintained its regulation, we considered its correlation with its nearest neighbor, in each species separately (score 1), as depicted in Figure 4E. Values lower than threshold indicate loss of regulation. To find whether the nearest neighbors of trans-diverging genes are regulatory-similar (Score 2), we calculated the correlation of a trans-diverging gene in a species A with the nearest neighbor of the same gene in species B. To avoid noise in the ranking of correlations, we considered the 5 nearest neighbors of species B and took the one that has the maximal correlation value with that gene in species A (Figure 4F). This was done similarly for *cis*-diverging genes by comparing hybrid alleles.

### Promoter analysis

Analyzed are sequences of S. cerevisiae and S. paradoxus are from Yue et al., 2017. Sequences of distant species were obtained from YGOB (Byrne and Wolfe, 2005). Motifs are from the YetFasco database (De Boer and Hughes, 2012).

## Acknowledgements

We thank the Barkai lab members and the Levy lab members for helpful discussions. We thank Patricia Wittkopp for commenting on the manuscript.

