## Supplementary material for "Independent evolution of transcript abundance and gene regulatory dynamics": Table S1, Table S2

### Supplementary materials:

Figures S1 to S5

Tables S1 and S2

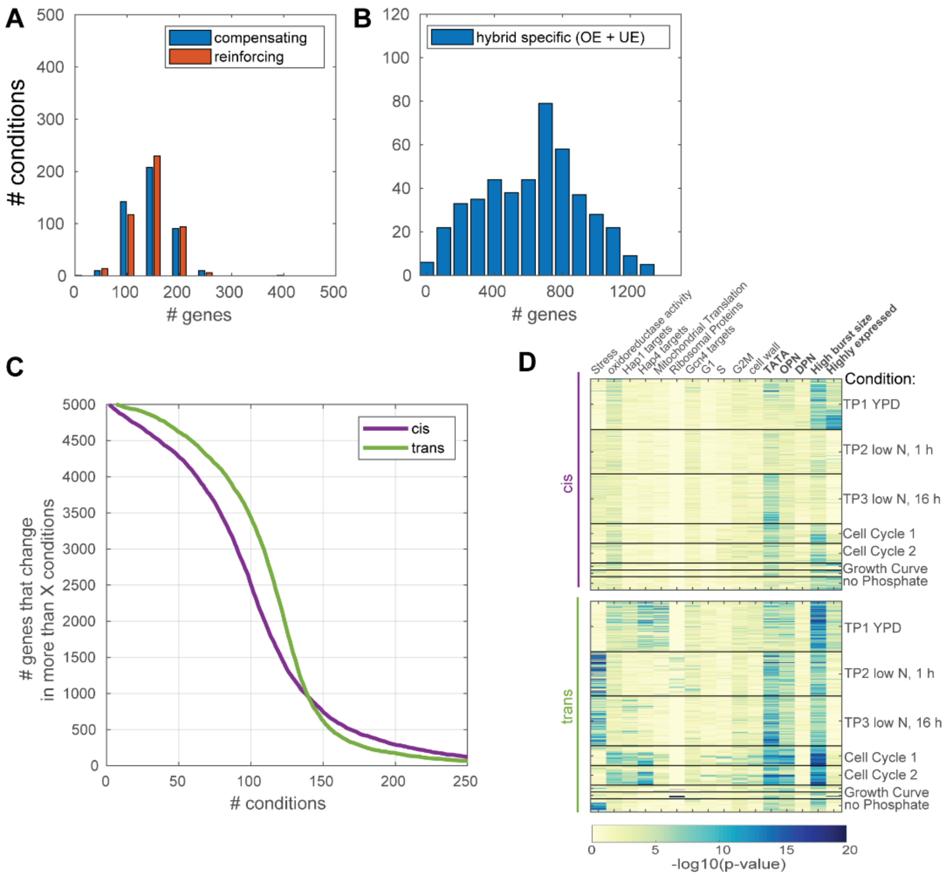


**Figure S 1:**

*A: compensating and reinforcing effects:* cis and trans effects can act one the same gene in different directions (compensating) or in the same direction (reinforcing). Shown here is the number of genes with cis- and trans-effects of >2 fold change, distinguished by the direction of the effect. Comparison of effects cross-conditions result in a similar number of compensating and reinforcing effects, as suggested in Fraser, 2019.

*B: Hybrid-specific effects:* Hybrid specific effects are cases where the hybrid over- or under-express a gene compared to both parents. Specifically here we considered the mean expression of the two hybrid alleles to be higher >2 folds from the maximal parental expression (over-expressed) or lower <2 folds from the minimal parental expression (under-expressed). We observe a median of 700 genes with hybrid-specific effects, using cross-condition comparison.

*C: cis-effects are more reproducible over conditions than trans-effects:* shown is an inverse cumulative histogram of the number of conditions a cis- or trans-effected gene showed a significance fold change (>2 folds, >1 z-score).

*D. functional gene groups are enriched in of cis- and trans-effected genes:* presented is the hypergeometric p-value for the indicated functional group with the cis- or trans-effected genes (>2 fold change), where each line represents a single condition.

**A**

| dataset name | publication | method | number of samples |
|---|---|---|---|
| Gasch | Gasch et al., 2000 | microarray | 134 |
| Kemmeren | Kemmeren et al., 2014 | microarray | 1487 |
| SPELL | Hibbs et al., 2007 | microarray | 7175 |
| Chapal | Chapal et al., 2019 | RNAseq | 2089 |
| Voichek | Voichek et al., 2018 | RNAseq | 838 |
| ours: cerevisiae | this study | RNAseq | 530 |
| ours: paradoxus | this study | RNAseq | 576 |
| ours: hyb cer | this study | RNAseq | 530 |
| ours:hyb par | this study | RNAseq | 530 |

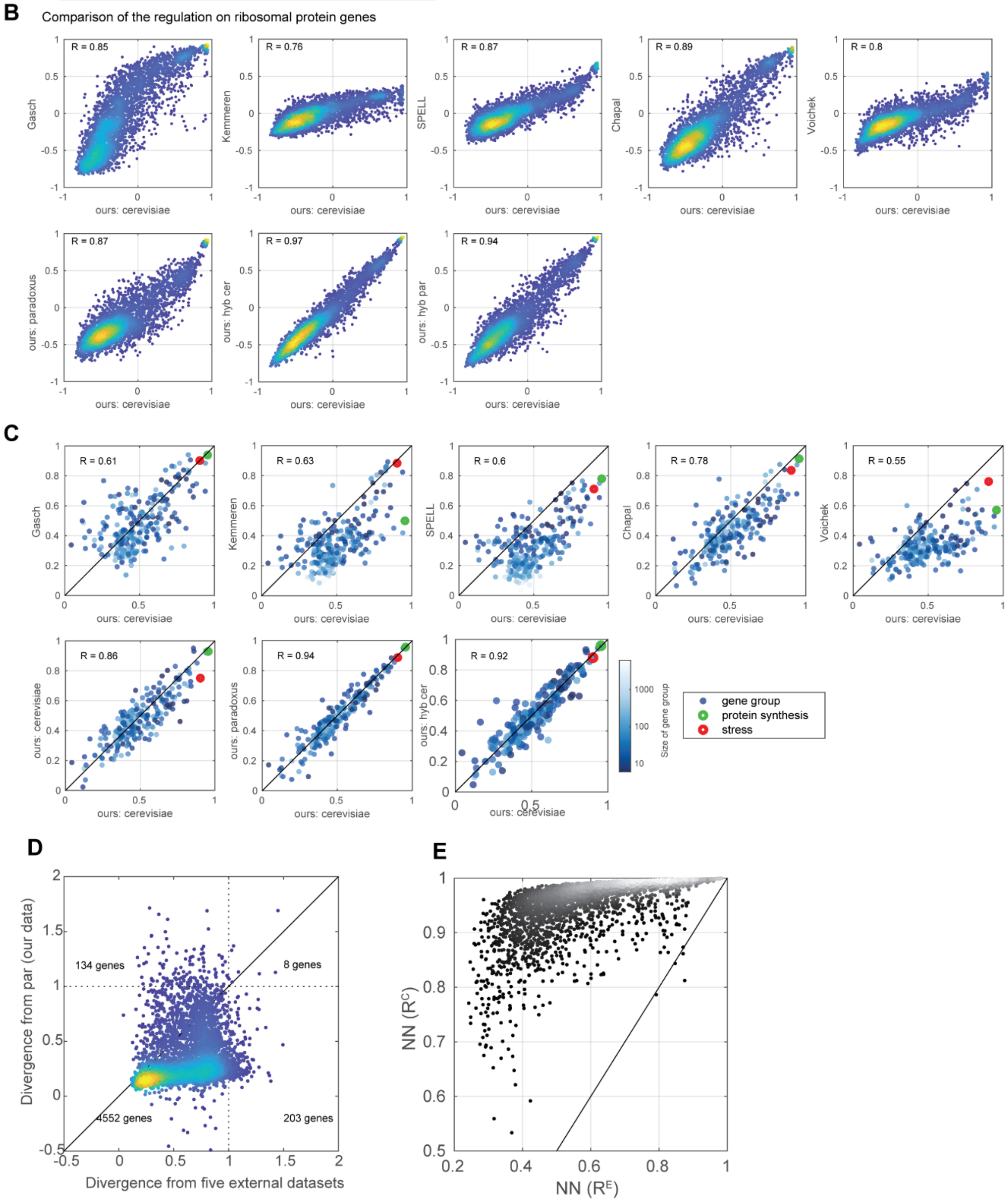

**Figure S 2:**

*A-D: Regulatory similarity of published expression datasets:*
*A: datasets used in the following comparisons* (Chapal et al., 2019; Gasch et al., 2000; Hibbs et al., 2007; Kemmeren et al., 2014; Voichek et al., 2018)
*B: Regulatory similarity of ribosomal protein genes:* same as in Figure 2B, the median correlation vector of 152 ribosomal protein genes was extracted for each dataset, and compared with S. cerevisiae dataset from this study.
*C: Consistency of functional gene groups:* Here other functional groups were considered (255 groups, see methods), where the mean correlation between genes in the group is presented, per dataset.
*D: Regulatory similarity per gene:* To measure difference in regulatory patterns between datasets we considered the dataset control. We normalize the similarity between two datasets (observed similarity) to the mean dataset control score (expected similarity), a measure which we term here divergence. Shown is the divergence of S. cerevisiae dataset in this study from five external dataset, compared with its divergence from S. paradoxus dataset of this study. Genes that diverge >1 from external datasets were filtered out from downstream analysis.

*E. Nearest-neighbor similarities for $R^E$ and $R^C$:* Shown are correlation-based similarities ($R^C$) between NNs, as a function expression-based similarities, $R^E$, computed from cerevisiae dataset.

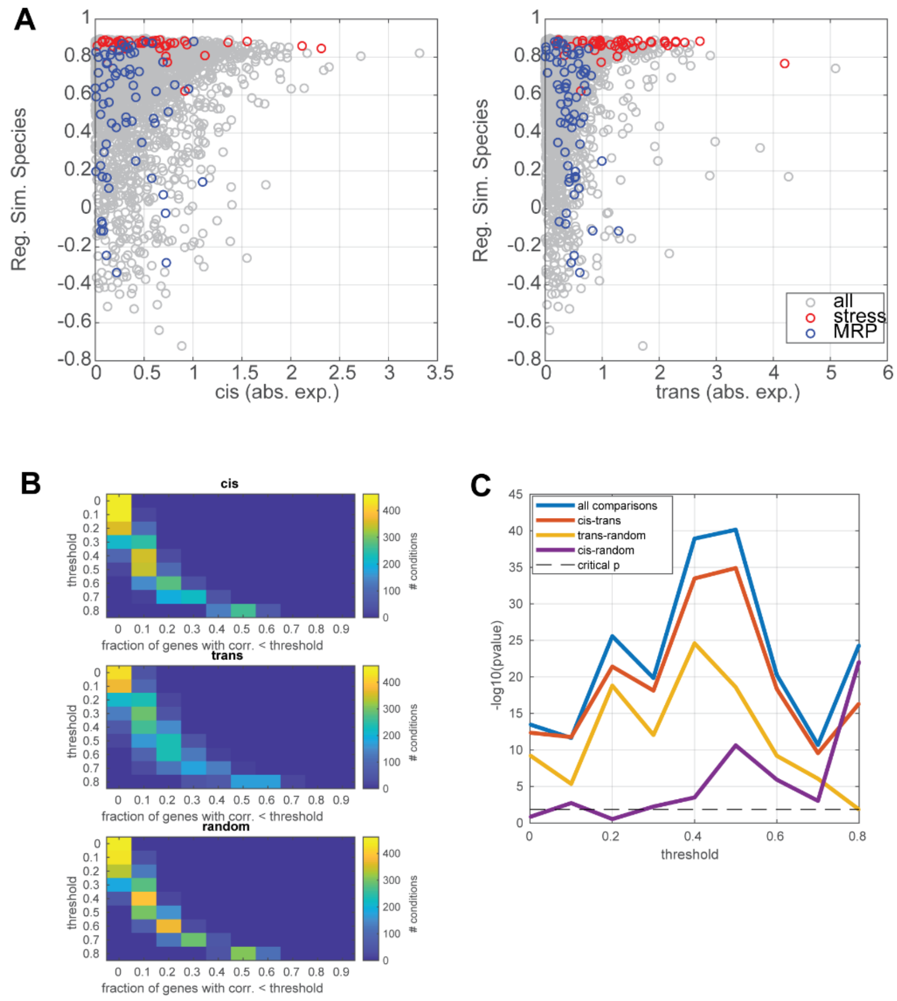


**Figure S 3:**

*A. Changes in absolute expression levels are independent of changes in regulation:* same as in Figure 3A, indicated groups are the stress and mitochondrial ribosomal protein modules.

*B-C. trans-dependent variations in absolute expression show some tendency for low regulatory similarity:* The measure used in Figure 3B is repeated for increasing thresholds of regulatory similarity. Shown are heatmaps of the distributions per genes (cis-effected, trans-effected or random set), per threshold (B), and the p-value (Kruskal-Wallis test) of the indicated comparison (C).

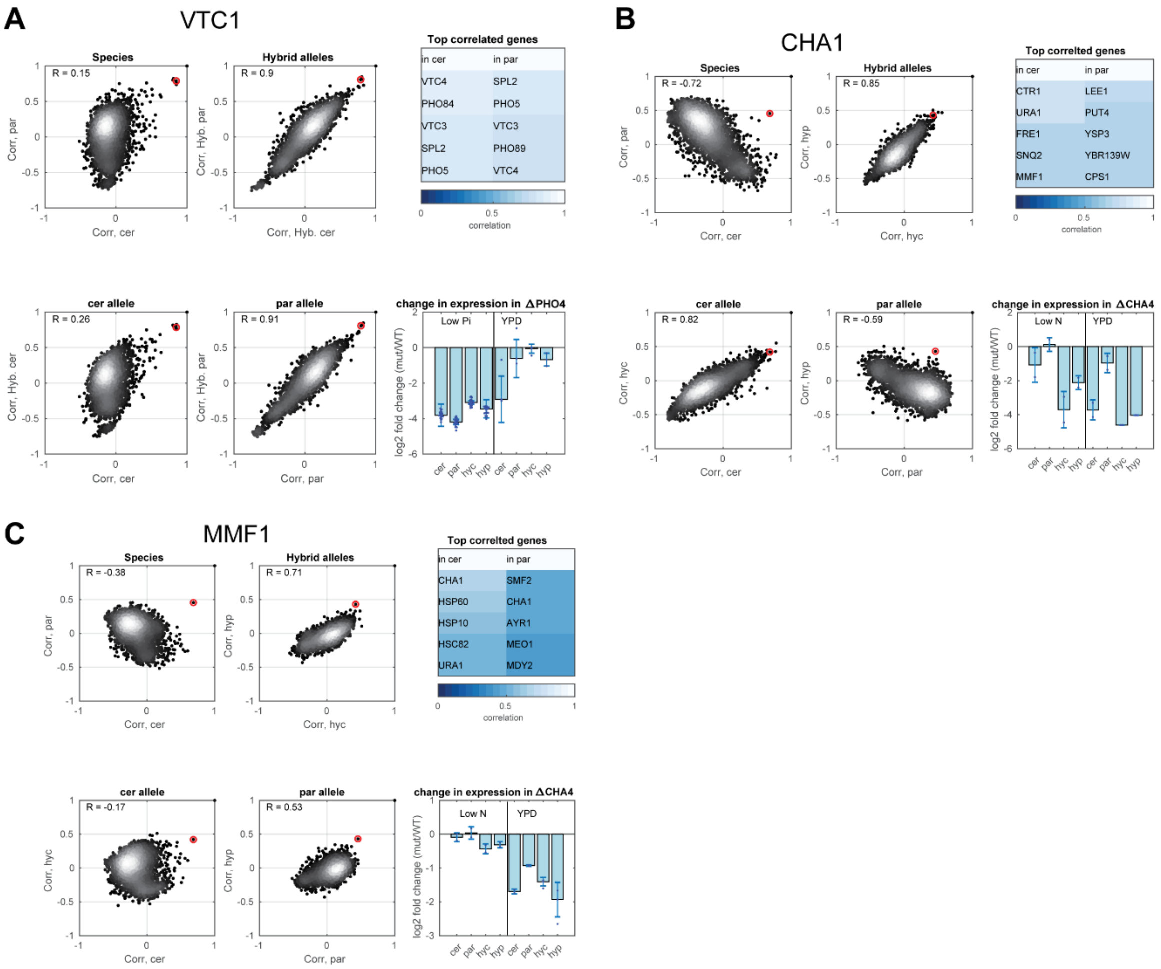


**Figure S 4:**

*A-C. nearest neighbors of trans-varying genes of which the direct-regulator is species-conserved:* shown as in Figure 4A, for VTC1 (regulated by Pho4, A) and for CHA1 and MMF1 (regulated by Cha4, B and C).

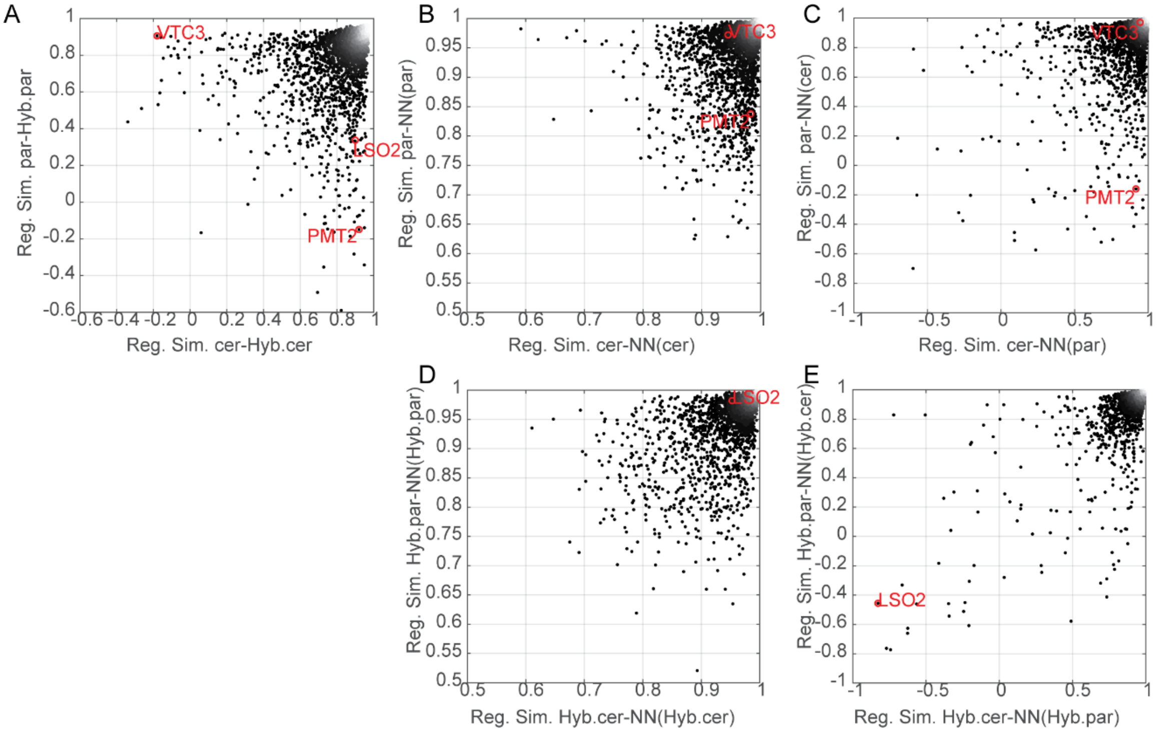


**Figure S 5:**

A. Regulatory similarity between each orthologue and its corresponding-allele in the hybrid: as in Figure 4C, shown for all figures. VTC3 and PMT2 are trans-varying, LSO2 is varies in cis.

B. nearest-neighbors similarities in the two orthologues. As in Figure 4D, shown for all genes.

C. cross-species nearest neighbor similarities. As in Figure 4E, shown for all genes.

D-E. same as in Figure 5 D-E, shown for all genes, indicated is the cis-varying gene LSO2.

Table S1: list of transcription factors deleted in the study. DBD family annotations are from De Boer and Hughes, 2012.

| no. | gene | orf name | Family (DBD) |
|---|---|---|---|
| 1 | ACE2 | YLR131C | C2H2 ZF |
| 2 | FKH1 | YIL131C | Forkhead |
| 3 | MBP1 | YDL056W | APSES |
| 4 | SKN7 | YHR206W | HSF |
| 5 | SWI4 | YER111C | APSES |
| 6 | SWI5 | YDR146C | C2H2 ZF |
| 7 | PHD1 | YKL043W | APSES |
| 8 | SOK2 | YMR016C | APSES |
| 9 | SUM1 | YDR310C | AT hook |
| 10 | TEC1 | YBR083W | TEA |
| 11 | HAP1 | YLR256W | Zinc cluster |
| 12 | MSN2 | YMR037C | C2H2 ZF |
| 13 | RFX1 | YLR176C | RFX |
| 14 | RIM101 | YHL027W | C2H2 ZF |
| 15 | ROX1 | YPR065W | Sox |
| 16 | STB5 | YHR178W | Zinc cluster |
| 17 | YAP1 | YML007W | bZIP |
| 18 | ZAP1 | YJL056C | C2H2 ZF |
| 19 | ARG80 | YMR042W | MADS box |
| 20 | ARG81 | YML099C | Zinc cluster |
| 21 | BAS1 | YKR099W | Myb/SANT |
| 22 | CBF1 | YJR060W | bHLH |
| 23 | GCN4 | YEL009C | bZIP |
| 24 | LEU3 | YLR451W | Zinc cluster |
| 25 | CHA4 | YLR098C | Zinc cluster |
| 26 | MIG1 | YGL035C | C2H2 ZF |
| 27 | NRG1 | YDR043C | C2H2 ZF |
| 28 | RGT1 | YKL038W | Zinc cluster |
| 29 | GCR2 | YNL199C | GCR1 |
| 30 | AFT1 | YGL071W | AFT |
| 31 | INO2 | YDR123C | bHLH |
| 32 | INO4 | YOL108C | bHLH |
| 33 | GLN3 | YER040W | GATA |
| 34 | GZF3 | YJL110C | GATA |
| 35 | PHO4 | YFR034C | bHLH |
| 36 | RPN4 | YDL020C | C2H2 ZF |
| 37 | HAP2 | YGL237C | |

| | | | |
|---|---|---|---|
| **38** | HAP3 | YBL021C | |
| **39** | HAP4 | YKL109W | CBF/NF-Y |
| **40** | HAP5 | YOR358W | |
| **41** | RTG1 | YOL067C | bHLH |
| **42** | RTG3 | YBL103C | bHLH |
| **43** | URE2 | YNL229C | |
| **44** | GAT1 | YFL021W | GATA |
| **45** | DAL80 | YKR034W | GATA |
| **46** | MET28 | YIR017C | |

Table S2: list of strains used in the study

| | strain | genotype |
|---|---|---|
| 1 | BY4741 | MATa his3Δ1 leu2Δ0 met15Δ0 ura3Δ0 |
| 2 | BY4742 | MATα his3Δ1 leu2Δ0 lys2Δ0 ura3Δ0 |
| 3 | S. cerevisiae a GFP WT | BY4741 HO::NatR-GFP |
| 4 | S. cereivisiae alpha GFP WT | BY4742 HO::HygR-GFP |
| 5 | S. cerevisiae diploid | strain 3 x strain 4 |
| 6 | S. paradoxus a mCherry WT | OS142 MATa HO::NatR-mCherry |
| 7 | S. paradoxus alpha mCherry WT | OS142 MATα ho::HygR-mCherry |
| 8 | S. paradoxus diploid | strain 3 x strain 4 |
| 9 | Hybrid | strain 3 x strain 6 |
| 10 | Hybrid in cell-cycle experiments | strain 3 x strain 6 |
| 11 | S. cerevisiae a ΔTFX | strain 3 TFXΔ::KanR |
| 12 | S. cerevisiae alpha ΔTFX | strain 4 TFXΔ::KanR |
| 13 | S. cereivisae diploid | strain 11 x strain 12 |
| 14 | S. paradoxus a ΔTFX | strain 6 TFXΔ::KanR |
| 15 | S. paradoxus alpha ΔTFX | strain 7 TFXΔ::KanR |
| 16 | S. paradoxus a ΔTFX | strain 14 x strain 15 |
| 17 | Hybrid ΔTFX | strain 11 x strain 15 |
